# A Comprehensive Survey of Genomic Mutations in Breast Cancer Reveals Recurrent Neoantigens as Potential Therapeutic Targets

**DOI:** 10.1101/2020.02.11.943258

**Authors:** Songming Liu, Chao Chen, Xiuqing Zhang

## Abstract

Neoantigens are newly formed antigens generated by cancer cells but absent in normal cells. With their high specificity and immunogenicity characteristic, neoantigens are considered as an ideal target for immunotherapy. This study was aimed to investigate the signature of neoantigens in breast cancer. Somatic mutations, including SNVs and indels, were obtained from cBioPortal of 5991 breast cancer patients. For neoantigen prediction, 738 non-silent somatic variants present in at least 3 patients were selected., PIK3CA (38%), the highly mutated gene in breast cancer, can produce the highest number of neoantigens per gene. Some pan-cancer hotspot mutations, such as PIK3CA E545K (6.93%), can be recognized by at least one HLA molecule. Since there are more SNVs than indels in breast cancer, SNVs are the major source of neoantigens. Patients with hormone receptor positive or HER2 positive are more competent to produce neoantigens. Age, but not clinical stage, is a significant contributory factor of neoantigens production. We believe a detailed description of breast cancer neoantigens signatures could contribute to neoantigens based immunotherapy development.

## 1. Introduction

Breast cancer is the world’s highest incidence of female malignant tumors, the incidence and mortality rate are ranked in the first female malignant tumors. There were more than two million new cases of breast cancer in 2018, with one in every four cancers of women diagnosed with the disease^[1]^. Breast cancer is a highly heterogeneous tumor that is currently classified by three molecular markers, including estrogen receptor (ER), progesterone receptor (PR) and HER2 (also called ERBB2), Treatment methods and prognosis of different subtypes of breast cancer vary greatly^[2, 3]^.

In recent years, cancer immunotherapy is playing an important role in a variety of solid tumors^[4-6]^. The most representative immunotherapy methods are immune checkpoint inhibitors (ICIs), including PD1/PD-L1 inhibitors, but ICIs are only about 30 percent effective. Neoantigen-based immunotherapy is a complement to ICIs therapy, such as neoantigen-based tumor-infiltrating lymphocytes (TILs) therapy in metastatic breast cancer^[7]^. However, the limitation of neoantigen-based immunotherapy is that different tumor patients have different mutations, and there are fewer neoantigens in common among different tumor patients. T-cell immunotherapy based on KRAS K12D mutation has been reported in colorectal cancer^[8]^, but similar therapies have not been reported in breast cancer.

With the development of sequencing technology, there have been many studies on the mutation characteristics of breast cancer, and a large number of articles based on second-generation sequencing technology have been published^[9-12]^. By integrating the clinical information and mutation data of the 8 previous breast cancer research cohorts., we obtained the mutation data of 5991 breast cancer patients^[3, 9-13]^. Finally, by combining the high-frequency HLA information and mutation data, we get the most common neoantigens in breast cancer patients, which provides a new road for the neoantigen-based immunotherapy.

## 2. Results and Discussion

### 2.1 The Mutation Landscape of Breast Cancer Patients

We have shown the mutation status of all samples in Figure 1 and Figure S1. Obviously, missense mutation is the main variant classification. At base substitution level, the dominant mutant form is C>T (Figure S2). Besides, the mutation load of each sample is pretty low, with only 25.3 mutations each sample on average and 6 mutations for the median. *PIK3CA* (38%) and *TP53* (37%) are two significantly mutated genes, with frequencies much higher than others, such as *GATA3* (12%) and *CDH1* (12%) (Figure 1F). Previous studies have described this high occurrence of *PIK3CA* and *TP53* mutations in breast cancer, although according to which the frequency of *TP53* is higher^[2, 3, 10, 14, 15]^.

**Figure 1.**
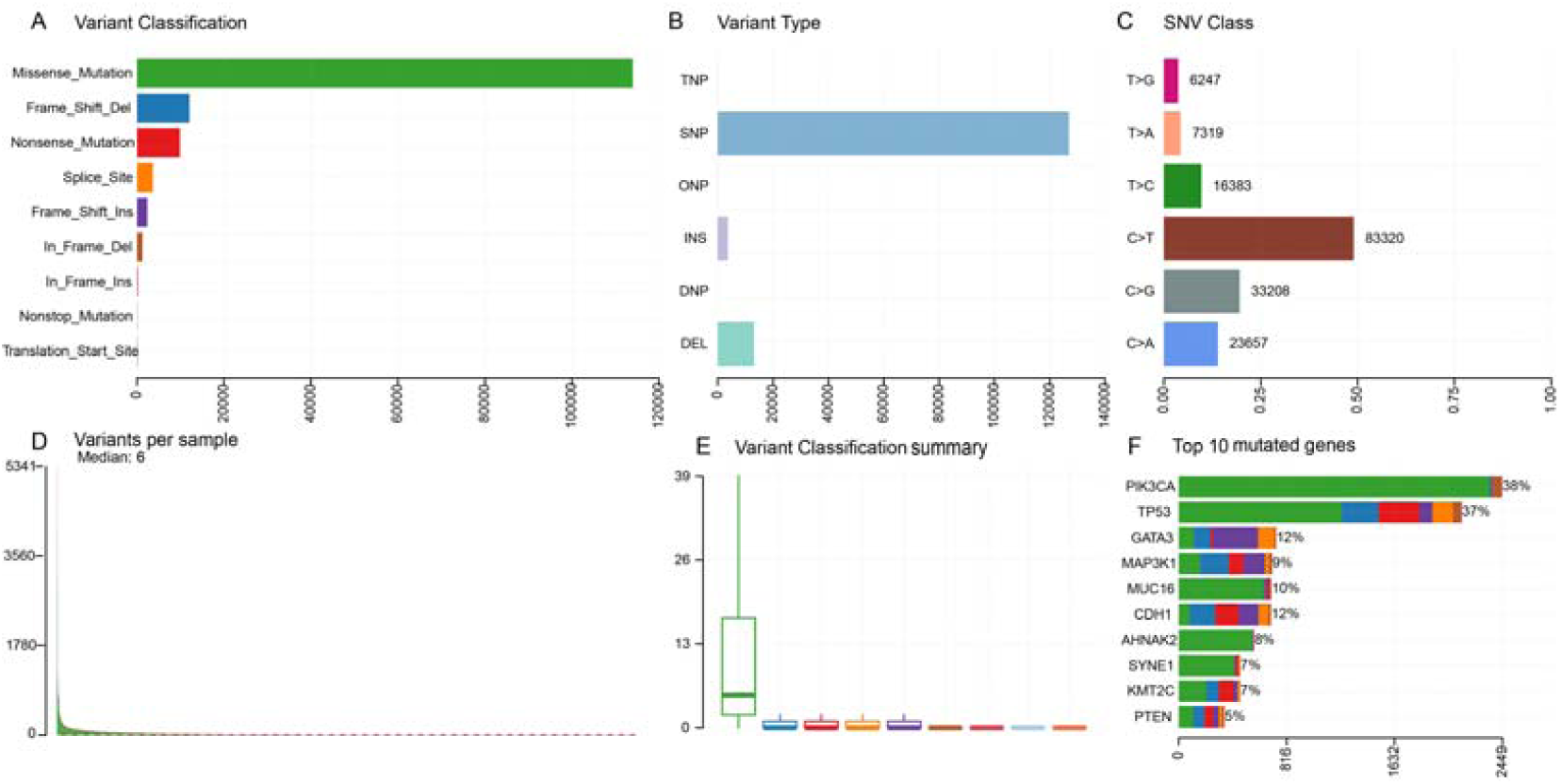
The mutation landscape of breast cancer cohort. **A**: Counts of each variant classification; **B:** Counts of each variant type; **C:** Counts of each SNV class; **D:** Mutation load per sample; **E**: Boxplot of each variant classification; **F**: Top 10 significantly mutant genes.

*TP53* and *PIK3CA* mutations occur frequently no matter in HER2+ or HER2-patients. However, the mutated rates are different, for which *TP53* (66%) is higher than *PIK3CA* (32%) in HER2+ patients but conversely in HER2-patients, with mutated rates 33% and 40%, respectively (Figure S3-S4). This may indicate its importance on the development and heterogeneity of breast cancer. In further investigation, we identified 22 significantly mutated genes between these two subgroups (Fisher’s exact test, *P* < 0.01, Figure S5). A similar result was observed between samples with hormone receptor-positive (HR+) status, defined as PR+ and/or ER+, and HR-status, defined as PR- and ER- (Figure S6-S8).

There are 895 recurrent mutations present in at least 3 patients, comprising 774 SNVs, 120 indels and 1 oligo-nucleotide polymorphism (ONP). Mutations after filtering silent variants, coupled with common HLA genotypes, were used for neoantigen prediction. Since previous studies have proved that the common neoantigens may serve as immunotherapy targets^[16, 17]^, we try to figure out whether there are common neoantigens in breast cancer populations in this way.

### 2.2 Results of Neoantigen Prediction

Due to the difference between HLA frequency of various populations, high-frequency (> 5%) HLA genotypes were selected from Han Chinese^[18]^ and Americans^[19]^ to predict “public” neoantigens (Table S1).

After filtering, there are 738 non-silent somatic variants, including 617 SNVs and 121 indels, with 356 and 86 derived peptides respectively (Table S2-S3). Not all variants exist corresponding neoantigens. Actually, SNV-related neoantigens derived from only 265 variants while 32 variants for indel-related neoantigens. In terms of SNVs, mutations of *PIK3CA, AKT1, SF3B1*, and *ESR1* produce the top 10 neoantigens with the highest frequency (Table 1), especially for *PIK3CA*, occupying 6 of 10. As for indels (Table 2), although the mutation frequency is lower, the number of neoantigens per mutation is higher, which is 2.69 for each indel but only 1.34 for each SNV on average.

**Table 1.** Top 10 SNVs and corresponding neoantigens.

| Chr | Location | Gene | AA change | Peptide | Frequency | HLA types |
| --- | --- | --- | --- | --- | --- | --- |
| chr3 | 178952085 | PIK3CA | H1047R | ARHGGWTTK | 839 | HLA-B27:05 |
| chr3 | 178936091 | PIK3CA | E545K | ITKQEKDFLW | 415 | HLA-B57:01 |
| chr3 | 178936091 | PIK3CA | E545K | STRDPLSEITK | 415 | HLA-A03:01; HLA-A11:01 |
| chr14 | 105246551 | AKT1 | E17K | RGKYIKTWR | 196 | HLA-A31:01 |
| chr3 | 178921553 | PIK3CA | N345K | ATYVKVNIR | 132 | HLA-A31:01 |
| chr2 | 198266834 | SF3B1 | K700E | QEVRTISAL | 83 | HLA-B40:01 |
| chr2 | 198266834 | SF3B1 | K700E | GLVDEQQEV | 83 | HLA-A02:01 |
| chr3 | 178938934 | PIK3CA | E726K | KTQKVQMKF | 64 | HLA-A32:01; HLA-B57:01 |
| chr6 | 152419926 | ESR1 | D538G | LYGLLLEML | 59 | HLA-A24:02 |
| chr3 | 178927980 | PIK3CA | C420R | KEEHRPLAW | 48 | HLA-B44:03 |

**Table 2.** Top 10 indels and corresponding neoantigens.

| Chr | Location | Gene | AA change | Peptide | Frequency | HLA types |
| --- | --- | --- | --- | --- | --- | --- |
| chr3 | 178916938 | PIK3CA | E110del | KVIEPVGNREK | 11 | HLA-A03:01; HLA-A11:01 |
| chr5 | 56177011 | MAP3K1 | R763Cfs*35 | LMFHKLSL | 8 | HLA-B08:01 |
| chr5 | 56177011 | MAP3K1 | R763Cfs*35 | FLLNFILIL | 8 | HLA-A02:01; HLA-A02:07 |
| chr5 | 56177011 | MAP3K1 | R763Cfs*35 | LILSVLMFH | 8 | HLA-A03:01 |
| chr5 | 56177011 | MAP3K1 | R763Cfs*35 | NFLLNFILI | 8 | HLA-A24:02 |
| chr5 | 56155721 | MAP3K1 | R273Sfs*27 | KSFPSAFSEW | 7 | HLA-B57:01 |
| chr5 | 56155721 | MAP3K1 | R273Sfs*27 | TTPKSPFTR | 7 | HLA-A11:01; HLA-A68:01 |
| chr5 | 67591104 | PIK3R1 | K567_L570del | KRMNSIIQLR | 7 | HLA-B27:05 |
| chr5 | 56155721 | MAP3K1 | R273Sfs*27 | SPFTRWLL | 7 | HLA-B08:01 |
| chr5 | 67591104 | PIK3R1 | K567_L570del | RMNSIIQLR | 7 | HLA-A31:01 |

### 2.3 Comparison of Neoantigens in Different Subgroups

To investigate the relations between clinical information and neoantigens, patients were divided into different subgroups by several clinical indexes. By comparing neoantigens in different subgroups, we found that the elder population (age > 60) carries more SNV-derived neoantigens (Fisher’s exact test, *P* < 0.01, Figure 2A). As for results of PR or ER status subgroups, patients of positive status tend to produce more SNV-derived neoantigens (Fisher’s exact test, *P* < 0.01, Figure 2C-D) than negative ones. On the contrary, neoantigens from SNVs are more produced by HER2-patients (Figure 2B).

**Figure 2.**
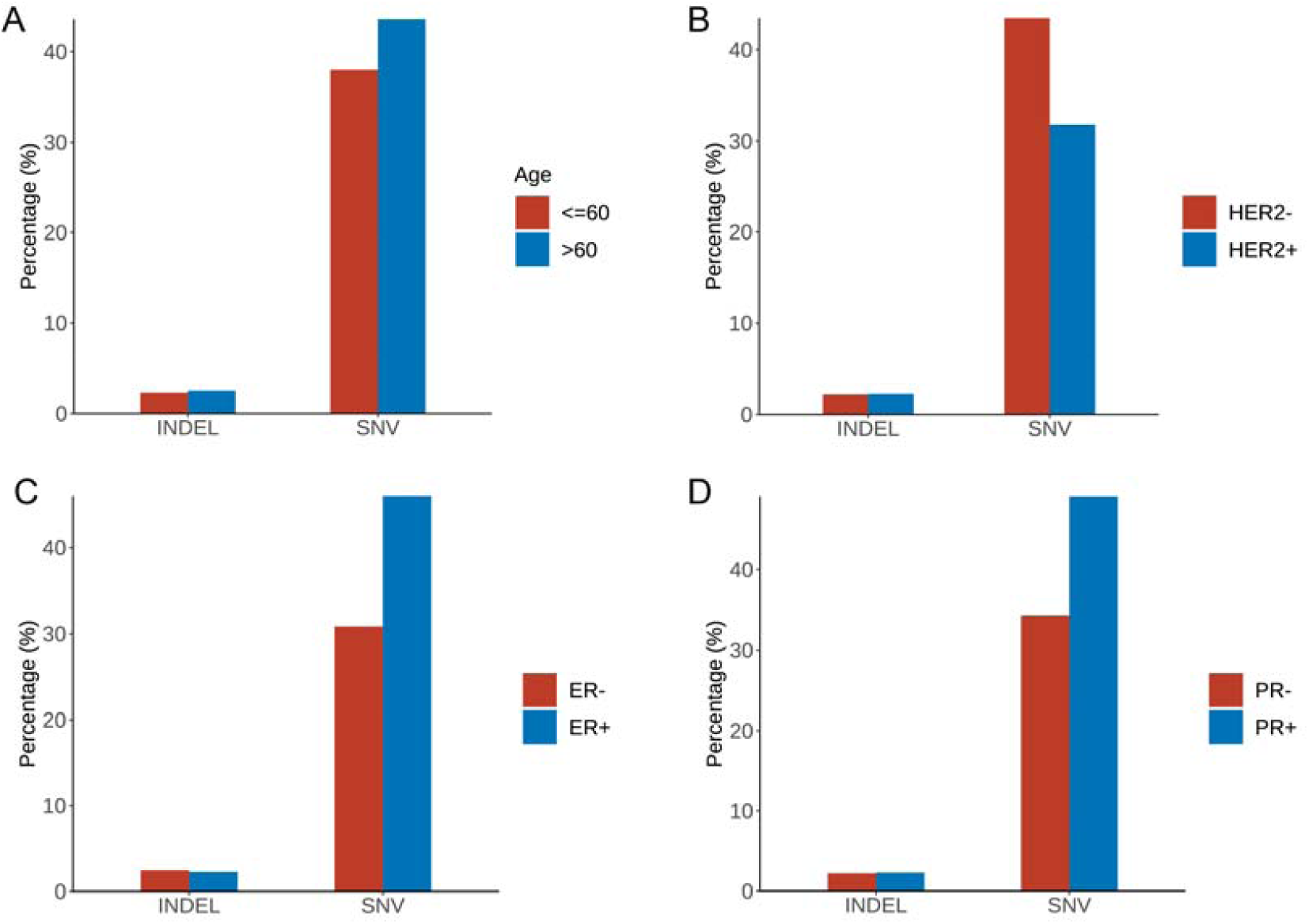
The comparison of neoantigens between different subgroups of breast cancer. **A**: Group Age: <=60 vs >60; **B**: Group HER2 status: HER2+ vs HER2-; **C**: Group ER status: ER+ vs ER-; **D**: Group PR status: PR+ vs PR-.

It seems that there is no significant difference in patients in different stages. Although we have observed the difference between patients in Stage I and Stage III (Fisher’s exact test, *P* < 0.01, Figure S9), no difference in other stages has reached the significance level. Notably, no matter in which subgroups, SNV-derived neoantigens can cover more patients than indes (Figure S10), suggesting that SNV is the major source of neoantigens in breast cancer patients.

### 2.4 Hotspot Mutations Derived Neoantigens May Serve as Targets of Immunotherapy in Breast Cancer and Pan-cancer

H1047R (*PIK3CA*), E545K (*PIK3CA*), E17K (*AKT1*), and N345K (*PIK3CA*) can produce recurrent neoantigens and have a higher mutation frequency in breast cancer cohort (Figure 3). Thus, we focused on these mutations to investigate the importance of themselves and corresponding neoantigens.

**Figure 3.**
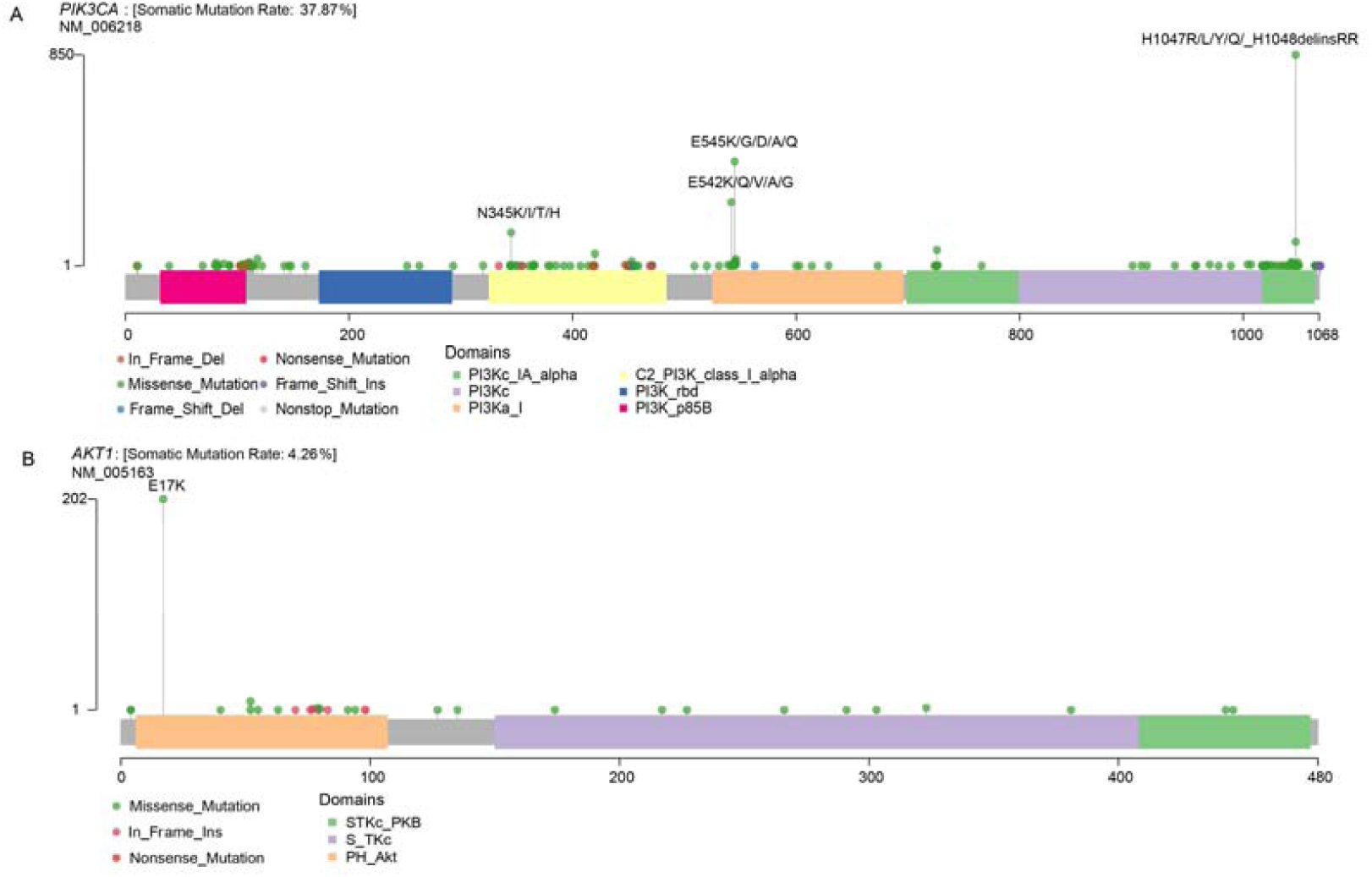
Mutational spectrum of specific genes. **A:** Mutations across the *PIK3CA* gene, no corresponding neoantigen for E542K; **B**: Mutations across the *AKT1* gene.

In this study, *PIK3CA* H1047R occurs in 14% of patients in our breast cancer cohort. This is consistent with the result of research by Zehir et al^[20]^. Besides, Meyer and colleagues demonstrate that this mutation in the luminal mammary epithelium can induce tumorigenesis^[21]^. In many other cancer types, this mutation also shows a pretty high frequency (Figure 4A). The previous study has shown H1047R can be presented by multiple HLA genotypes, such as HLA-C*07:02 and HLA-C*07:01^[16]^. These results imply that *PIK3CA* H1047R may be used as a source of “public” neoantigen in many cancer types.

**Figure 4.**
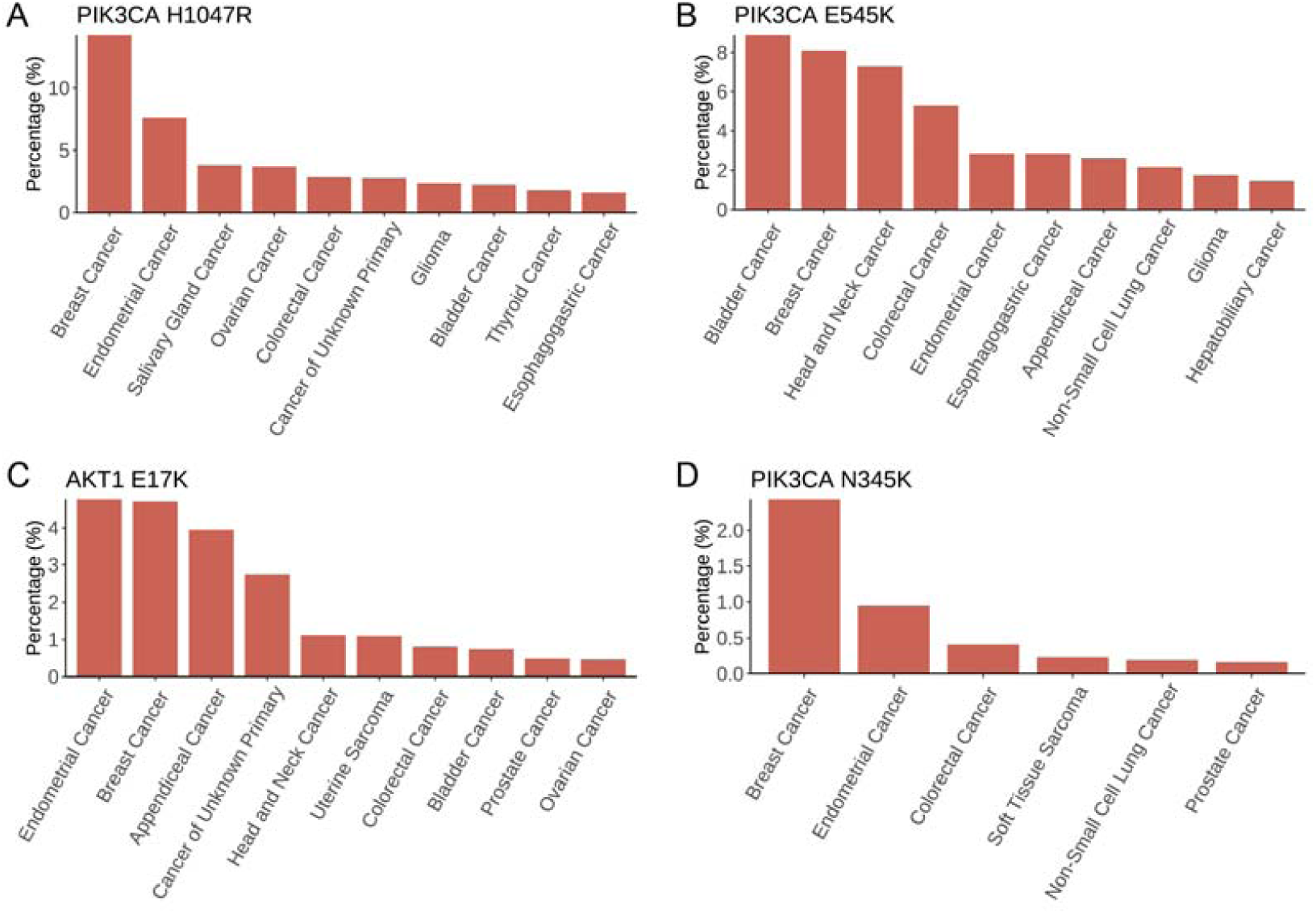
Mutation frequency across multiple cancer types in MSK-IMPACT cohorts. **A**: *PIK3CA* H1047R; **B**: *PIK3CA* E545K; **C:** *AKT1* E17K; **D:** *PIK3CA* N345K. All cancer types shown above should meet the following criteria: 1) with a total number of patients equals or exceed 50 in MSK-IMPACT cohort; 2) mutated frequency of the corresponding mutation should exceed 0; 3) if there are over 10 cancer types, show only the top 10 results with the highest frequency.

*PIK3CA* E545K is a hotspot mutation with first-line drugs^[22]^. In breast cancer, this mutation shows a frequency of about 8%, second to bladder cancer (Figure 4B). As for *PIK3CA* N345K, its mutation frequency is pretty low across all cancers shown in Figure 4D. A case report suggests this mutation might be associated with the sensitivity of Everolimus^[23]^.

*AKT1* E17K occurs in many solid tumors with a low frequency (Figure 4C). Compared to other *AKT1* mutations, E17K shows a much higher occurrence (Figure 3B). In certain breast cancer patients, this mutation is the most likely disease driver^[24]^. Subsequently, a study demonstrates *AKT1* E17K is a therapeutic target in many cancers^[25]^.

## 3. Conclusion

Based on the analysis of mutation data from eight breast cancer studies, we described the most complete mutation landscape of breast cancer so far. 43 HLA genotypes with high frequency in Chinese or TCGA cohort and 738 non-silent somatic mutations were selected to predict the common neoantigens. The high-frequency mutations, including *PIK3CA* H1047R (14%), *PIK3CA* E545K (6.93%), *AKT1* E17K (3.27%) and *PIK3CA* N345K (2.20%), can be recognized by multiple HLA molecules, such as HLA-A1101 and HLA-A03:01. These HLA genotypes are the dominant HLA subtypes in the Han Chinese and Americans, representing the commonality of neoantigens we identified among breast cancer patients. In conclusion, except for having constructed a comprehensive mutation landscape of breast cancer, we also have found a serious of public neoantigens, which may contribute to the development of immunotherapy in breast cancer.

## 4. Materials and Methods

### 4.1 Genomic Data for Breast Cancer Patients

All somatic mutations, including single nucleotide variants (SNVs) and short insertion/deletion (indels), were obtained from the published datasets. The data comprises 5991 breast cancer patients from eight studies, covering several important studies, such as The Cancer Genome Atlas Program (TCGA). Clinical information is showed in Table 3 and Table S4. There is no need for additional informed consent cause all data were from public databases with informed consent provided in the original studies.

**Table 3.** Summary of clinical information of 5991 patients from eight studies.

| Characteristic | BCCRC<br>(n=65) | BROAD<br>(n=103) | IGR<br>(n=216) | MBCproject<br>(n=180) | METABRIC<br>(n=2509) | MSK<br>(n=1746) | Sanger<br>(n=100) | TCGA<br>(n=1072) | Total<br>(n=5991) |
| --- | --- | --- | --- | --- | --- | --- | --- | --- | --- |
| **Age (years)** |  |  |  |  |  |  |  |  |  |
| <=60 | 41 (63.1%) | 86 (83.5%) | 0 (0%) | 107 (59.4%) | 1163 (46.4%) | 1333 (76.3%) | 65 (65.0%) | 593 (55.3%) | 3388 (56.6%) |
| >60 | 22 (33.8%) | 17 (16.5%) | 0 (0%) | 8 (4.4%) | 1335 (53.2%) | 413 (23.7%) | 35 (35.0%) | 479 (44.7%) | 2309 (38.5%) |
| Unknown | 2 (3.1%) | 0 (0%) | 216 (100%) | 65 (36.1%) | 11 (0.4%) | 0 (0%) | 0 (0%) | 0 (0%) | 294 (4.9%) |
| **Stage** |  |  |  |  |  |  |  |  |  |
| 0 | 0 (0%) | 0 (0%) | 0 (0%) | 0 (0%) | 24 (1.0%) | 0 (0%) | 0 (0%) | 0 (0%) | 24 (0.4%) |
| I | 0 (0%) | 11 (10.7%) | 0 (0%) | 11 (6.1%) | 630 (25.1%) | 469 (26.9%) | 0 (0%) | 181 (16.9%) | 1302 (21.7%) |
| II | 0 (0%) | 73 (70.9%) | 0 (0%) | 27 (15.0%) | 979 (39.0%) | 512 (29.3%) | 0 (0%) | 608 (56.7%) | 2199 (36.7%) |
| III | 0 (0%) | 19 (18.4%) | 0 (0%) | 21 (11.7%) | 144 (5.7%) | 370 (21.2%) | 0 (0%) | 246 (22.9%) | 800 (13.4%) |
| IV | 0 (0%) | 0 (0%) | 0 (0%) | 51 (28.3%) | 11 (0.4%) | 381 (21.8%) | 0 (0%) | 18 (1.7%) | 461 (7.7%) |
| Unknown | 65 (100%) | 0 (0%) | 216 (100%) | 70 (38.9%) | 721 (28.7%) | 14 (0.8%) | 100 (100%) | 19 (1.8%) | 1205 (20.1%) |
| **ER Status** |  |  |  |  |  |  |  |  |  |
| Positive | 3 (4.6%) | 44 (42.7%) | 0 (0%) | 94 (52.2%) | 1825 (72.7%) | 1372 (78.6%) | 79 (79.0%) | 0 (0%) | 3417 (57.0%) |
| Negative | 61 (93.8%) | 28 (27.2%) | 0 (0%) | 19 (10.6%) | 644 (25.7%) | 329 (18.8%) | 21 (21.0%) | 0 (0%) | 1102 (18.4%) |
| Unknown | 1 (1.5%) | 31 (30.1%) | 216 (100%) | 67 (37.2%) | 40 (1.6%) | 45 (2.6%) | 0 (0%) | 1072 (100%) | 1472 (24.6%) |
| **PR Status** |  |  |  |  |  |  |  |  |  |
| Positive | 1 (1.5%) | 40 (38.8%) | 0 (0%) | 83 (46.1%) | 1040 (41.5%) | 994 (56.9%) | 60 (60.0%) | 0 (0%) | 2218 (37.0%) |
| Negative | 63 (96.9%) | 32 (31.1%) | 0 (0%) | 28 (15.6%) | 940 (37.5%) | 692 (39.6%) | 40 (40.0%) | 0 (0%) | 1795 (30.0%) |
| Unknown | 1 (1.5%) | 31 (30.1%) | 216 (100%) | 69 (38.3%) | 529 (21.1%) | 60 (3.4%) | 0 (0%) | 1072 (100%) | 1978 (33.0%) |
| **HER2 Status** |  |  |  |  |  |  |  |  |  |
| Positive | 0 (0%) | 8 (7.8%) | 14 (6.5%) | 37 (20.6%) | 247 (9.8%) | 145 (8.3%) | 30 (30.0%) | 0 (0%) | 481 (8.0%) |
| Negative | 63 (96.9%) | 47 (45.6%) | 194 (89.8%) | 71 (39.4%) | 1733 (69.1%) | 1248 (71.5%) | 70 (70.0%) | 0 (0%) | 3426 (57.2%) |
| Unknown | 2 (3.1%) | 48 (46.6%) | 8 (3.7%) | 72 (40.0%) | 529 (21.1%) | 353 (20.2%) | 0 (0%) | 1072 (100%) | 2084 (34.8%) |

### 4.2 Neoantigen Prediction

Mutations should be non-silent somatic mutations and identified in at least 3 breast cancer patients. Subsequently, These mutations, combining with 43 high-frequency HLA genotypes in Chinese and TCGA cohorts, were used to predict neoantigens through NetMHC^[26]^, NetMHCpan^[27]^, PSSMHCpan^[28]^, PickPocket^[29]^ and SMM^[30]^. Criteria for neoantigen screening refer to our previous research ^[16]^.

### 4.3 Statistical Analysis

We finished all statistical analyses in R-Studio (R version 3.6.0). The two R packages, maftools^[31]^ and ggplot2^[32]^, were used for mutations analysis and visualization, respectively. Unless special instruction was given, *P* < 0.01 was considered significant.

## Acknowledgements

This work was supported by the Science, Technology and Innovation Commission of Shenzhen Municipality under grant No. GJHZ20170314152701465.

